# Failed down-regulation of PI3K signaling makes autoreactive B cells receptive to bystander T cell help

**DOI:** 10.1101/2023.01.23.525206

**Authors:** Brigita E. Fiske, Andrew Getahun

## Abstract

The role of T cell help in autoantibody responses is not well understood. Since tolerance mechanisms govern both T and B cell responses, one might predict that both T cell tolerance and B cell tolerance must be defeated in autoantibody responses requiring T cell help. To define whether autoreactive B cells depend on T cells to generate autoantibody responses, we studied the role of T cells in autoantibody responses resulting from acute cell-specific deletion of regulatory phosphatases. Ars/A1 B cells are DNA-reactive and require continuous inhibitory signaling by the tyrosine phosphatase SHP-1 and the inositol phosphatases SHIP-1 and PTEN to maintain unresponsiveness. Acute B cell-restricted deletion of any of these phosphatases results in an autoantibody response. Here we show that CD40-CD40L interactions are required to support autoantibody responses of B cells whose anergy has been compromised. If the B cell-intrinsic driver of loss of tolerance is failed negative regulation of PI3K signaling, bystander T cells provide sufficient CD40-mediated signal 2 to support an autoantibody response. However, while autoantibody responses driven by acute B cell-targeted deletion of SHP-1 also require T cells, bystander T cell help does not suffice. These results demonstrate that upregulation of PI3K signaling in autoreactive B cells, recapitulating the effect of multiple autoimmunity risk alleles, promotes autoantibody responses both by increasing B cells’ cooperation with non-cognate T cell help, as well as by altering BCR signaling. Receptiveness to bystander T cell help enables autoreactive B cells to circumvent the fail-safe of T cell tolerance.

**Significance:** Phosphatase suppression of PI3K signaling is an important mechanism by which peripheral autoreactive B cells are kept in an unresponsive/anergic state. Loss of this suppression, due to genetic alleles that confer risk of autoimmunity, often occurs in autoreactive B cells of individuals who develop autoimmune disease. Here we demonstrate that de-repression of PI3K signaling promotes autoantibody responses of a DNA-reactive B cell clone by relaxing dependence of autoantibody responses on T cell-derived helper signals. These results suggest that impaired regulation of PI3K signaling can promote autoantibody responses in two ways: by restoring antigen receptor signaling and by enabling autoreactive B cells to circumvent restrictions imposed by T cell tolerance mechanisms.

## Introduction

The two-signal hypothesis of lymphocyte activation has played a central role in our evolving understanding of the initiation of immune responses and immune tolerance (1). Receiving a signal through the antigen receptor (signal 1) causes an initial activation of lymphocytes. However, a second signal (signal 2) generally is required for productive immune responses. If lymphocytes receive signal 1 without signal 2, chronic antigen receptor signals become tolerogenic, and the lymphocytes acquire an unresponsive (anergic) state. For B cells, signal 2 is often provided by T cells. In the case of autoreactive B cells present in the periphery, T cell tolerance mechanisms would be expected to eliminate or limit the availability of (functional) antigen-specific T cells that could provide that second signal, providing an extra layer of tolerance to prevent autoantibody responses (2, 3).

A significant proportion of the autoreactive B cells present in the periphery display an anergic phenotype (4–7); these include B cells with autoantigen-specificity relevant for major autoimmune diseases such as diabetes (8), SLE (9) and rheumatoid arthritis (10, 11). Studies using B cell receptor transgenic models of anergy and studies that examine anergic B cells found within the wild type repertoire have revealed that anergic B cells exist on a continuum, with multiple mechanisms contributing to a degree of unresponsiveness (12–14). Early work using the MD4 x ML5 model of B cell anergy established that suppression of BCR signaling is the key determinant of anergic B cell unresponsiveness (15). Anergic B cells still may respond to non-BCR signals, but in general, these signals are insufficient to support an autoantibody response when BCR signaling (signal 1) is suppressed (15–17).

Several mechanisms limit the ability of anergic autoreactive B cells to receive sufficient signaling through their antigen receptors to support lymphocyte activation, including downregulation of surface IgM expression and active inhibitory signaling (12–14). Anergic B cells interact continuously with self-antigen through their antigen receptors, and this continuous antigen receptor engagement and signaling, possibly combined with co-ligation of inhibitory receptors, causes increased activity of regulatory phosphatases. These include inositol lipid phosphatases such as PTEN and SHIP-1 that suppress the PI3K pathway and tyrosine phosphatases such as SHP-1 that can dephosphorylate key tyrosine residues on early actors in the BCR signaling cascade (18–20).

In an earlier study, we examined the requirement for continuous inhibitory signaling to maintain B cell unresponsiveness of anergic B cells using the Ars/A1 model. Ars/A1 B cells express a transgenic BCR reactive with the hapten *p*-azophenylarsonate (Ars), which also binds DNA-containing self-antigens, rendering the B cells anergic (21). We used an adoptive transfer approach in which we transferred mature dilution dye-loaded anergic Ars/A1 B cells into recipient mice and induced deletion of relevant phosphatases in the transferred B cells using an inducible cre-lox system. We found that both continuous suppression of the PI3K pathway by SHIP-1 and continuous suppressive activity by SHP-1 are required to maintain unresponsiveness (19). Deletion of either phosphatase results in restoration of BCR signaling and autoantigen-driven proliferation and differentiation into autoantibody secreting plasmablasts.

Conditions sufficient to drive autoimmunity in this experimental system are B cell intrinsic loss of SHIP-1, PTEN or SHP-1, derepressing antigen receptor signaling. DNA-reactive B cells, such as Ars/A1 B cells, respond to self-antigens containing Toll-like receptor (TLR) ligands which could, in theory, provide a second signal needed for lymphocyte activation by activating TLRs (22). Thus, upon loss of effector phosphatases, Ars/A1 B cells receive all signals needed to mount an autoantibody response. Here we assessed the role of B cell extrinsic signals provided by T cells in autoantibody responses by anergy-compromised Ars/A1 B cells. We found that while B cell intrinsic loss of phosphatase activity allows B cell activation, CD40-CD40L interaction-mediated signal 2 is required for autoantibody responses. Importantly, when B cell-intrinsic dysregulation of PI3K signaling is the driver of loss of tolerance, bystander T cells are able to provide sufficient CD40L-mediated help to support an autoantibody response, effectively enabling the B cells to circumvent restrictions imposed by T cell tolerance.

## Results

### Compromised Ars/A1 B cells require CD40L-mediated signals to mount an autoantibody response

Many autoantibody responses are either dependent upon or augmented by T cell help and CD40-CD40L dependent B-T cell interactions (23–26). These include responses by autoreactive B cells that recognize self-antigens that contain TLR ligands, such as DNA, where theoretically the TLR could provide signal 2. To interrogate the role of T cell help mediated by CD40-CD40L interactions in autoantibody responses of SHIP-1 deficient Ars/A1 B cells, we tested the ability of anti-CD40L to block responses. We used a previously described adoptive transfer model (19) (Fig 1A) where mature anergic Ars/A1 B cells are loaded with a dilution dye and adoptively transferred into low-dose irradiated recipient mice. These Ars/A1 B cells express an inducible Cre recombinase and have a YFP reporter with a stop-flox-cassette knocked into the Rosa26 locus, in addition to a floxed SHIP-1 gene. After the adoptive transfer, the recipient mice are treated with tamoxifen to induce Cre activation in the transferred Ars/A1 B cells, generating a population of YFP+ SHIP-1 deficient Ars/A1 B cells. In control mice, Ars/A1 B cells were transferred that express the inducible Cre recombinase, have the YFP reporter construct, but have a WT SHIP-1 locus instead of the floxed SHIP-1 locus. Tamoxifen treatment results in the generation of YFP+ SHIP-1 sufficient Ars/A1 B cells (WT Ars/A1) that remain anergic. As described previously (19), 14 days after tamoxifen treatment SHIP-1 deficient Ars/A1 B cells (SHIP-1 Ars/A1) found in the spleen have proliferated (Fig 1B-C, isotype plots) and differentiated into plasmablasts, as evident by CD138 expression (Fig 1B-C, isotype plots) and numeration of antibody secreting cells in the spleen (Fig 1D, isotype plots). In contrast, Ars/A1 B cells that remain SHIP-1 sufficient (WT Ars/A1) did not proliferate or differentiate into plasmablasts (Fig 1B-D, isotype plots). Blocking CD40-CD40L interactions with anti-CD40L blocking antibody clone MR-1 completely prevented the accumulation of proliferated and differentiated SHIP-1-deficient Ars/A1 B cells and resultant autoantibody production (Fig 1B-D, anti-CD40L plots). In this situation, the absence of proliferated and differentiated SHIP-1 deficient Ars/A1 B cells could be due to an absence of a response or due to a survival defect in responding SHIP-1 deficient Ars/A1 B cells. To distinguish between these two possibilities, we quantified the number of undivided and total YFP+ Ars/A1 B cells in anti-CD40L- or isotype-treated mice. While we observed a trend towards a reduction in YFP+ SHIP-1 sufficient Ars/A1 B cells (Ars WT) in MR-1 treated mice (Fig 1E), there was not a significant difference in two independent experiments. In contrast, there was a stark reduction in the number of undivided and total SHIP-1 deficient YPP+ Ars/A1 B cells in anti-CD40L treated mice compared to isotype treated mice (Fig 1E). This suggests that the lack of accumulation of proliferated and differentiated YFP+ SHIP-1 deficient Ars/A1 B cells in anti-CD40L-treated mice is due to a lack of CD40-mediated survival signals, preventing a form of activation-induced cell death, rather than the lack of a CD40-mediated activation signal that is required for the SHIP-1 deficient Ars/A1 B cell to respond.

**Figure 1.**
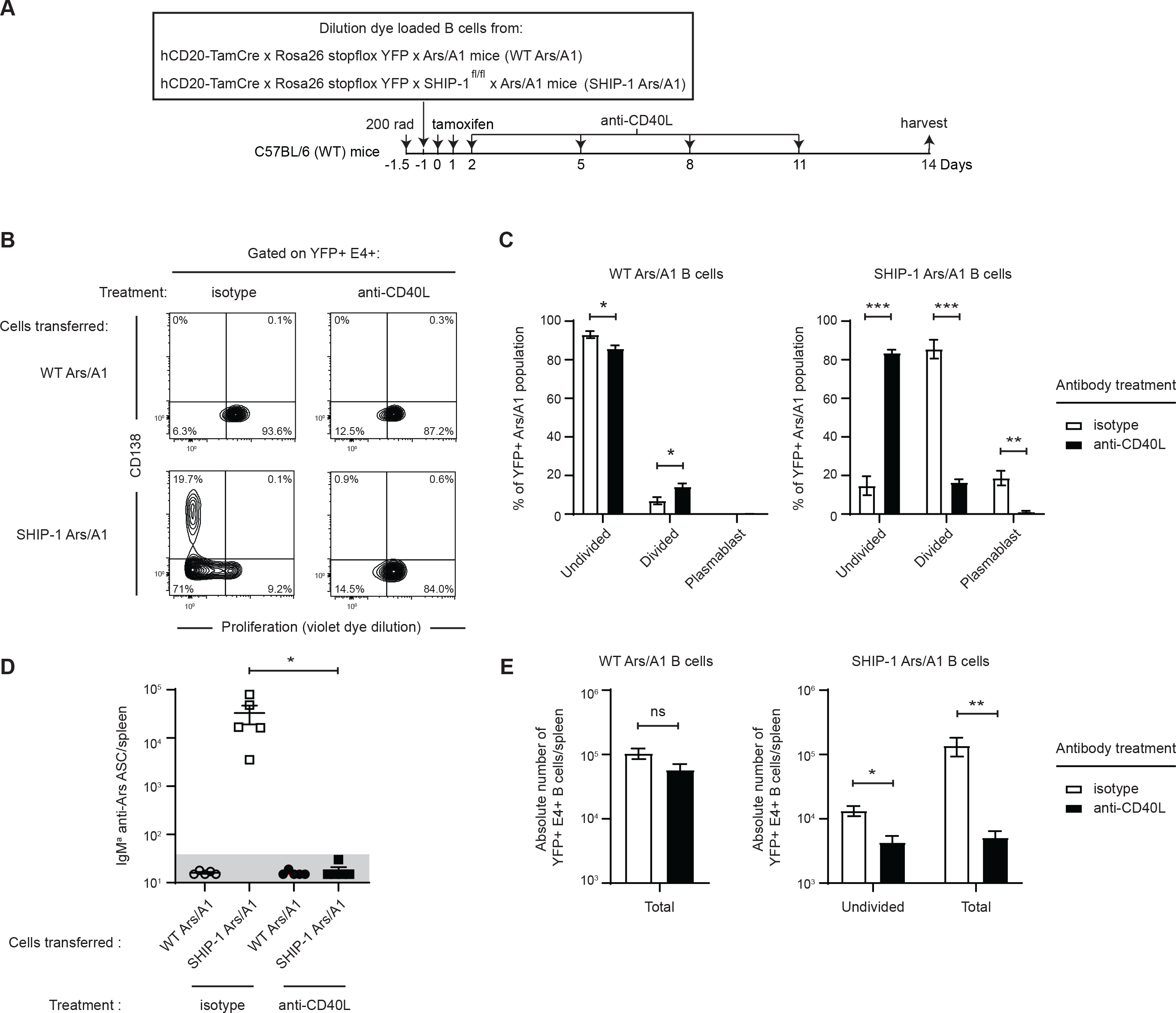
SHIP-1 deficient Ars/A1 B cells require CD40L-mediated signals to mount an autoantibody response. A) Schematic representation of the experimental protocol. Recipient mice received anti-CD40L or isotype control antibody on the indicated days. B) Proliferation and differentiation to plasmablasts of splenic YFP+ E4+ SHIP-1 sufficient (WT Ars/A1) and SHIP-1 deficient (SHIP Ars/A1) B cells 14 days after tamoxifen treatment (n=5/group, representative cytograms shown). C) Distribution of the YFP+ E4+ B cell population shown in Fig 1B between an undivided, proliferated and plasmablast (proliferated and CD138+) state. D) Quantification of antibody secreting cells (IgM^a^ anti-Ars) by ELISPOT of spleen cells from Fig 1B. Gray area delineates the limit of detection (50 spots/spleen). E) Quantification of the number of YFP+ E4+ cells/spleen in the indicated transfers 14 days after tamoxifen treatment. Data shown are representative of at least two replicate experiments. Error bars represent mean ± SEM. Two-tailed unpaired Student’s *t* test was used. ns, P > 0.05; *, P < 0.05; **, P < 0.01; ***, P < 0.001.

### CD4 T cells facilitate autoantibody responses of SHIP-1-compromised Ars/A1 B cells

CD4 T cells are the predominant source of CD40L signals for antibody responses. To determine whether compromised Ars/A1 B cell responses require CD4 T cells, we adoptively transferred SHIP-1 Ars/A1 B cells into WT or TCRα KO mice and assayed their response 14 days following induction of SHIP-1 deletion (Fig 2A). At this timepoint, the recipient TCRα KO mice had <1% CD4+ T cells in the splenic lymphocyte population (data not shown). The near absence of CD4 T cells in recipient TCRα KO mice severely impaired the accumulation of proliferated and differentiated YFP+ SHIP-1 deficient B cells (Fig 2 B-C) and resultant generation of autoantibody secreting cells (Fig 2D). Since TCRα KO mice lack both CD4 and CD8 T cells, we adoptively transferred B cells into CD8 KO recipient mice to exclude the possibility of CD8 T cell involvement. Deficiency of CD8 T cells in recipient mice did not impair the ability of SHIP-1 deficient Ars/A1 B cells to mount an autoantibody response (Fig 2 E-G). These results suggest that CD4 T cells play an important role in providing CD40L-derived help to compromised Ars/A1 B cells.

**Figure 2.**
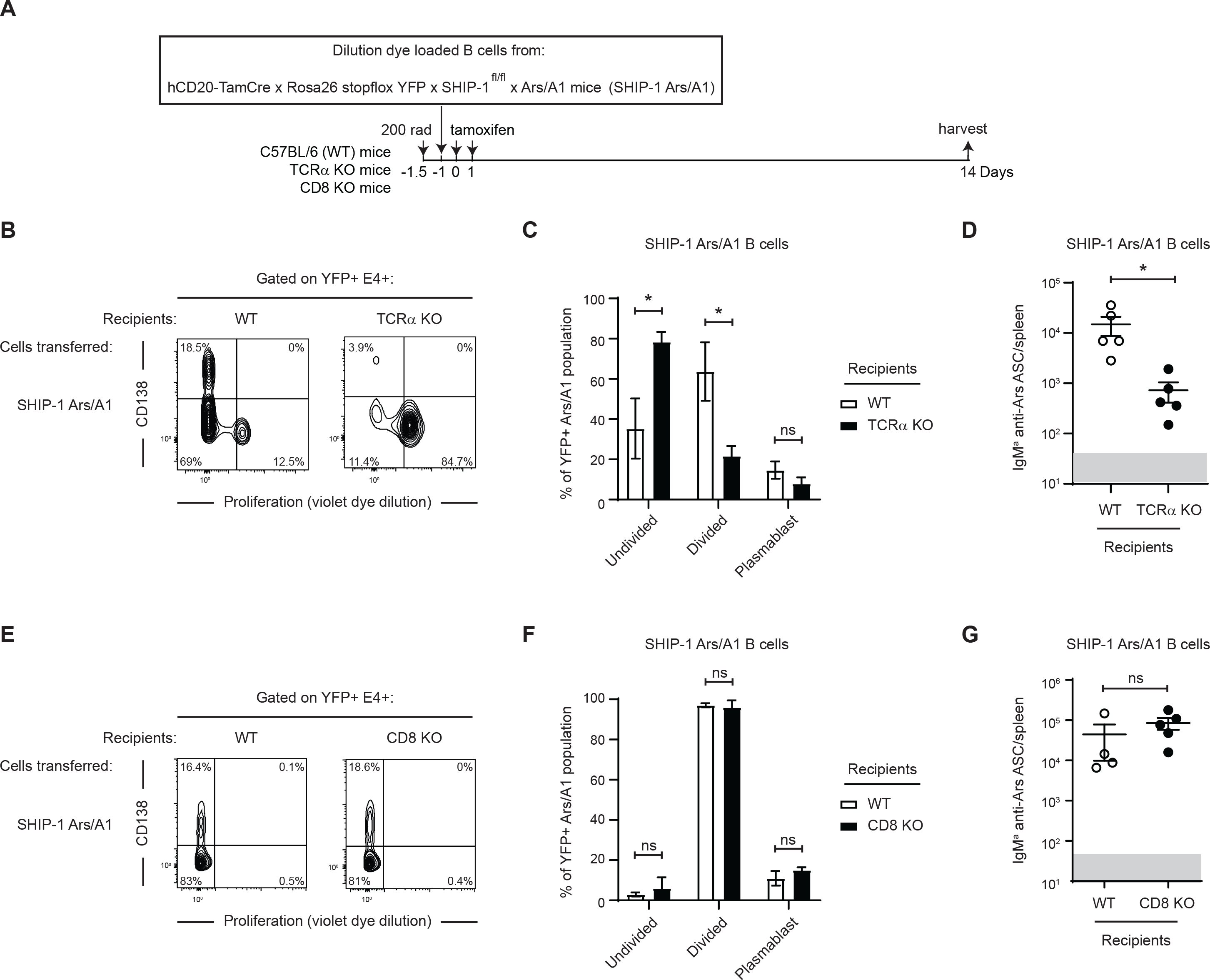
CD4 T cells facilitate autoantibody responses of SHIP-1 deficient Ars/A1 B cells. A) Schematic representation of the experimental protocol. In the adoptive transfer experiments into TCRα KO mice, the recipient mice received an injection with a depleting anti-TCRβ antibody (clone H57-597) on days 3 and 7 post transfer to deplete any T cells that were transferred into these mice during the B cell transfer. B) Proliferation and differentiation to plasmablasts of splenic YFP+ E4+ SHIP-1 deficient B cells in WT C57BL/6 mice and TCRα KO mice 14 days after tamoxifen treatment (n=5/group, representative cytograms shown). C) Distribution of the YFP+ E4+ SHIP-1 deficient Ars/A1 B cell population shown in Fig 2B between an undivided, proliferated and plasmablast (proliferated and CD138+) state. D) Quantification of antibody secreting cells (IgM^a^ anti-Ars) by ELISPOT of spleen cells from Fig 2B. Gray area delineates the limit of detection (50 spots/spleen). E) Proliferation and differentiation to plasmablasts of splenic YFP+ E4+ SHIP-1 deficient B cells in WT C57BL/6 mice and CD8 KO mice 14 days after tamoxifen treatment (n=4-5/group, representative cytograms shown). F) Distribution of the YFP+ E4+ SHIP-1 deficient Ars/A1 B cell population shown in Fig 2E between an undivided, proliferated and plasmablast (proliferated and CD138+) state. G) Quantification of antibody secreting cells (IgM^a^ anti-Ars) by ELISPOT of spleen cells from Fig 2E. Gray area delineates the limit of detection (50 spots/spleen). Data shown are representative of at least two replicate experiments. Error bars represent mean ± SEM. Two-tailed unpaired Student’s *t* test was used. ns, P > 0.05; *, P < 0.05; **, P < 0.01; ***, P < 0.001.

### Bystander CD4 T cells can support autoantibody responses of SHIP-1 deficient Ars/A1 B cells

Next, we determined if the CD4 T cells needed to be autoantigen-reactive to support autoantibody responses. We crossed OTII TCR transgenic mice onto the TCRα^−/−^ background to generate mice that only have OVA-specific CD4 T cells and used these mice as recipients (Fig 3A). Interestingly, the presence of non-cognate CD4 T cells was sufficient to support autoantibody responses by SHIP-1 deficient Ars/A1 B cells as measured by the number of autoantibody secreting Ars/A1 B cells/spleen (Fig 3D). Analysis of the phenotype of YFP+ SHIP-1 deficient Ars/A1 B cells revealed that while the frequency of proliferated CD138+ Ars/A1 B cells was comparable to WT recipients with a full CD4 T cell repertoire, the frequency of proliferated SHIP-1 deficient Ars/A1 B cells was significantly reduced in the WT recipients (Fig 3B-C). These results indicate that bystander CD4 T cells are sufficient to support compromised Ars/A1 B cells, although cognate T cell help is required for an optimal B cell response.

**Figure 3.**
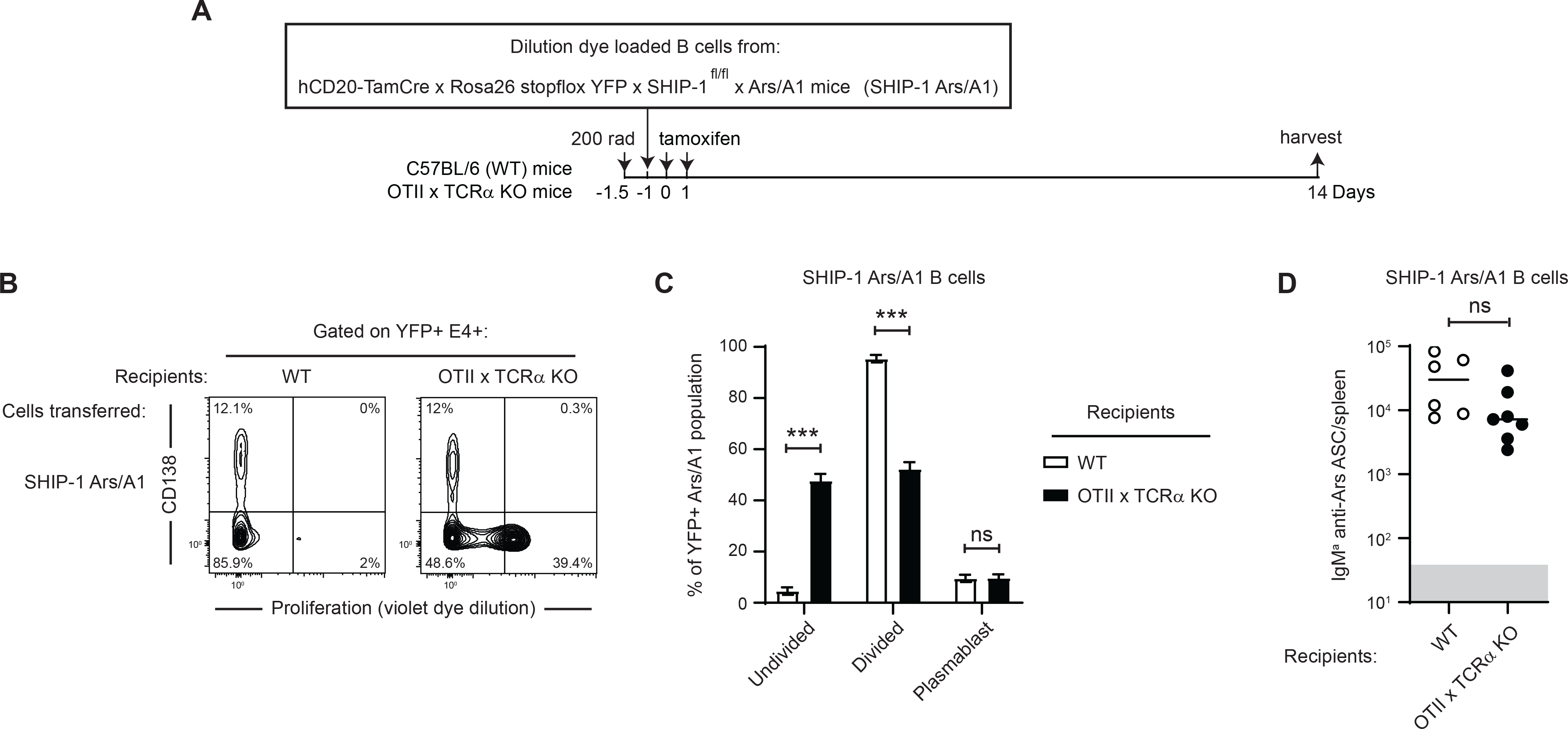
Bystander CD4 T cells can support autoantibody responses of SHIP-1 deficient Ars/A1 B cells. A) Schematic representation of the experimental protocol. B) Proliferation and differentiation to plasmablasts of splenic YFP+ E4+ SHIP-1 deficient B cells in WT C57BL/6 mice and OTII x TCRα KO mice 14 days after tamoxifen treatment (n=6-7/group, representative cytograms shown). C) Distribution of the YFP+ E4+ B cell population shown in Fig 3B between an undivided, proliferated and plasmablast (proliferated and CD138+) state. D) Quantification of antibody secreting cells (IgM^a^ anti-Ars) by ELISPOT of spleen cells from Fig 3B. Gray area delineates the limit of detection (50 spots/spleen). Data shown are representative of at least two replicate experiments. Error bars represent mean ± SEM. Two-tailed unpaired Student’s *t* test was used. ns, P > 0.05; *, P < 0.05; **, P < 0.01; ***, P < 0.001.

### Cooperation of compromised Ars/A1 B cells and bystander CD4 T cell help is dependent on the nature of the B cell-intrinsic driver of the loss of tolerance

The data thus far indicate that bystander T cells can provide CD40 signals required for SHIP-1 deficient Ars/A1 B cells to mount an autoantibody response. To determine if this cooperative response occurs uniquely when anergy is compromised by SHIP-1 deficiency, we tested the cooperativity of bystander T cell help and Ars/A1 B cells in which anergy was compromised by different drivers (Fig 4A). First, we tested whether Ars/A1 B cells that were compromised by loss of another negative regulator of the PI3K pathway, the inositol phosphatase PTEN, are receptive to bystander T cell help. As previously described (19), induced loss of PTEN in Ars/A1 B cells following adoptive transfer into WT C57BL/6 mice led to proliferation and differentiation of YFP+ PTEN-deficient Ars/A1 B cells and to autoantibody forming cell generation (Fig 4B-D). When transferred into OTII x TCRα KO recipients, induced loss of PTEN in Ars/A1 B cells resulted in a response comparable to that observed in mice with a full CD4 T cell repertoire (Fig 4B-D). Next, we tested whether the cooperation with bystander T cells occurs when Ars/A1 B cells are compromised by loss of the tyrosine phosphatase SHP-1. SHP-1 dephosphorylates tyrosine residues on early actors in the BCR signaling cascade. While SHP-1 deficient Ars/A1 B cells proliferated and differentiated into autoantibody secreting cells in WT recipients (Fig 4E-G), their response was impaired in OTII x TCRα KO recipients (Fig 4 E-G). Enumeration of CD45.1+ E4+ B cells in spleens showed that few SHP-1 deficient Ars/A1 B cells survived in OTII x TCRα KO recipients and those that did did not proliferate (Fig 4H). These results suggest that the receptiveness of compromised Ars/A1 B cells to bystander T cell help is not a trait of the Ars/A1 clone *per se*, but rather depends on the B cell intrinsic change that breaks tolerance. B cell intrinsic hyperactivity of PI3K signaling increases the cooperativity of B cells with bystander T cell help. This could be due to a more advanced state of B cell activation or could reflect a role for PI3K in CD40 signaling.

**Figure 4.**
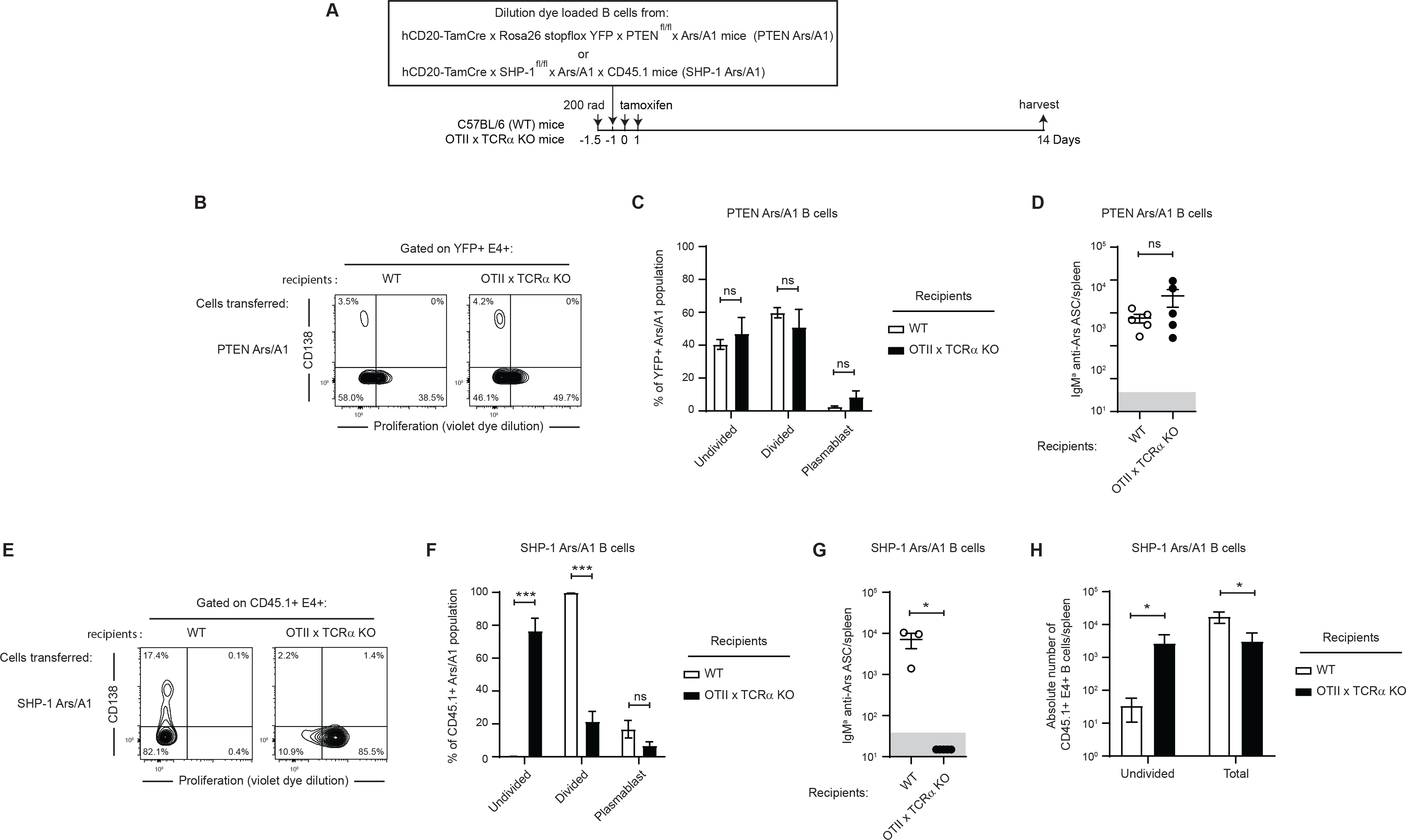
Bystander CD4 T cells can support autoantibody responses of PTEN deficient Ars/A1 B cells but not SHP-1 deficient Ars/A1 B cells. A) Schematic representation of the experimental protocol. B) Proliferation and differentiation to plasmablasts of splenic YFP+ E4+ PTEN deficient B cells in WT C57BL/6 mice and OTII x TCRα KO mice 14 days after tamoxifen treatment (n=5/group, representative cytograms shown). C) Distribution of the YFP+ E4+ PTEN deficient Ars/A1 B cell population shown in Fig 4B between an undivided, proliferated and plasmablast (proliferated and CD138+) state. D) Quantification of antibody secreting cells (IgM^a^ anti-Ars) by ELISPOT of spleen cells from Fig 4B. Gray area delineates the limit of detection (50 spots/spleen). E) Proliferation and differentiation to plasmablasts of splenic CD45.1+ E4+ SHP-1 deficient B cells in WT C57BL/6 mice and OTII x TCRα KO mice 14 days after tamoxifen treatment (n=3-5/group, representative cytograms shown). F) Distribution of the CD45.1+ E4+ SHP-1 deficient Ars/A1 B cell population shown in Fig 4E between an undivided, proliferated and plasmablast (proliferated and CD138+) state. G) Quantification of antibody secreting cells (IgM^a^ anti-Ars) by ELISPOT of spleen cells from Fig 4E. Gray area delineates the limit of detection (50 spots/spleen). H) Quantification of the number of CD45.1+ E4+ cells/spleen in the indicated transfers 14 days after tamoxifen treatment. Data shown are representative of at least two replicate experiments. Error bars represent mean ± SEM. Two-tailed unpaired Student’s *t* test was used. ns, P > 0.05; *, P < 0.05; **, P < 0.01; ***, P < 0.001.

### CD40 signals alone do not overcome B cell anergy

Compromised Ars/A1 B cells depend on CD40L-derived signals to mount an autoantibody response (Fig 1). The absence of an autoantibody response by SHP-1 deficient Ars/A1 B cells when only bystander T cell help is available (Fig 4E-H) suggests that CD40L-CD40 interactions are limiting under these conditions. To determine if CD40-derived signals were limiting or if additional signals were missing, we used an agonistic anti-CD40 antibody to supplement CD40 signaling (Fig 5A). Administration of an agonistic anti-CD40 antibody rescued SHP-1 deficient Ars/A1 B cells in OTII x TCRα KO recipients from activation-induced cell death and restored the antibody response (Fig 5B-E). This suggests that these B cells did not receive sufficient CD40 signals to make an effective plasmablast response when only bystander T cell help was available. To exclude the possibility that administration of an agonistic anti-CD40 antibody per se is sufficient to drive the loss of tolerance of anergic Ars/A1 B cells, we did control experiments in which WT Ars/A1 B cells were transferred into C57BL/6 recipients and exposed to a similar regiment of agonistic anti-CD40 antibody treatment. While we observed some proliferation of WT Ars/A1 B cells upon exposure to agonistic anti-CD40 antibody, these B cells did not develop into autoantibody secreting cells (Fig 5F-H). These data suggest that while CD40 signals are required for the development of Ars/A1 plasmablast responses, CD40 signals alone are insufficient to support an autoantibody response by anergic Ars/A1 B cells. Only when antigen receptor signaling (signal 1) has been restored do these autoreactive B cells become productively receptive to T cell derived signals.

**Figure 5.**
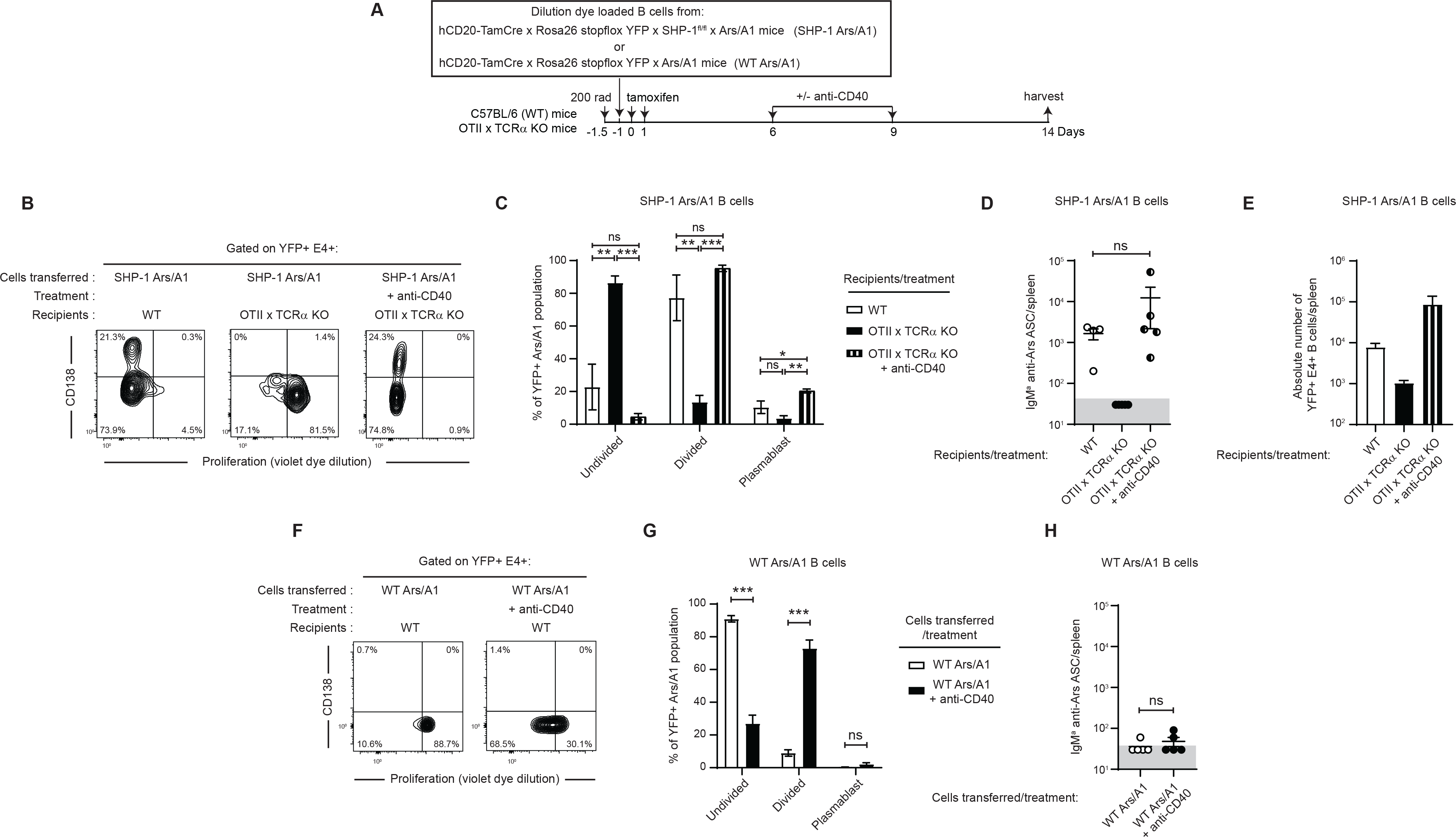
CD40 signals alone do not overcome B cell anergy. A) Schematic representation of the experimental protocol. One group of OTII x TCRα KO mice was treated with agonistic anti-CD40 antibody on days 6 and 9 post initiation of tamoxifen treatement. B) Proliferation and differentiation to plasmablasts of splenic YFP+ E4+ SHP-1 deficient B cells in WT C57BL/6 mice and OTII x TCRα KO mice, in the presence or absence of agonistic anti-CD40, 14 days after tamoxifen treatment (n=4-5/group, representative cytograms shown). C) Distribution of the YFP+ E4+ SHP-1 deficient Ars/A1 B cell population shown in Fig 5B between an undivided, proliferated and plasmablast (proliferated and CD138+) state. D) Quantification of antibody secreting cells (IgM^a^ anti-Ars) by ELISPOT of spleen cells from Fig 5B. Gray area delineates the limit of detection (50 spots/spleen). E) Quantification of the number of CD45.1+ E4+ cells/spleen in the indicated transfers 14 days after tamoxifen treatment. F) Proliferation and differentiation to plasmablasts of splenic YFP+ E4+ WT Ars/A1 B cells in WT C57BL/6 mice, in the presence or absence of agonistic anti-CD40, 14 days after tamoxifen treatment (n=5/group, representative cytograms shown). G) Distribution of the YFP+ E4+ WT Ars/A1 B cell population shown in Fig 5F between an undivided, proliferated and plasmablast (proliferated and CD138+) state. H) Quantification of antibody secreting cells (IgM^a^ anti-Ars) by ELISPOT of spleen cells from Fig 5F. Gray area delineates the limit of detection (50 spots/spleen). Data shown are representative of at least two replicate experiments. Error bars represent mean ± SEM. Two -tailed unpaired Student’s *t* test was used. ns, P > 0.05; *, P < 0.05; **, P < 0.01; ***, P < 0.001.

## Discussion

Here we show that while acute B cell intrinsic loss of tyrosine or inositol phosphatases is sufficient to cause a loss of tolerance of DNA/chromatin-reactive Ars/A1 B cells (19), B cell extrinsic factors, in this case, cognate or non-cognate T cell help, are necessary for this loss of B cell tolerance to lead to an autoantibody response. Specifically, we demonstrate that the auto-immune response of compromised Ars/A1 B cells requires CD40-mediated second signals. CD4 T cells are required for efficient autoantibody responses, suggesting that they suffice as a source of CD40L-induced CD40-mediated signals. Interestingly, whether these B-T cell interactions need to be cognate for this autoreactive B cell clone appears to depend on the B cell intrinsic changes that drive a loss of tolerance. Bystander T cells are sufficient to support autoantibody responses if the B cell intrinsic driver of loss of tolerance relaxes the regulation of PI3K signaling. Taken together, these data show that B-T cell interactions play an important role in the autoantibody response of compromised nuclear autoantigen-reactive B cells. Furthermore, the findings underscore the importance of appropriate regulation of PI3K signaling for maintenance of B cell tolerance and suggest a mechanism by which changes in PI3K signaling in autoreactive B cells may make them receptive to bystander T cell help, effectively enabling them to circumvent a layer of T cell tolerance mechanisms.

Ars/A1 B cell receptors bind DNA-containing self-antigens, including chromatin. These self-antigens contain TLR ligands, including TLR9 ligands. TLR9 can convey secondary signals that support B cell activation and plasma cell differentiation (22, 27). Therefore, antigen recognition by compromised Ars/A1 B cells should provide both signal 1 (via the BCR) and signal 2 (via TLR9) to Ars/A1 B cells, supporting autoantibody responses. In anergic B cells, including Ars/A1 B cells, efficient localization of BCR-bound DNA to TLR9-containing late endosomes is disrupted, interfering with TLR9 signaling (28). However, a reversal of anergy restores normal BCR/Ag trafficking and TLR9 signaling (28), indicating that in compromised Ars/A1 B cells BCR-associated TLR9 signaling will be restored as well. TLR9’s role in autoimmune disease has been a topic of debate. TLR9 is required for anti-DNA autoantibody responses (29, 30), yet in preclinical models, TLR9 appears to have a protective role while TLR7 has been identified as a promotor of autoimmune disease (30, 31). Recent work from the Cancro lab suggests a mechanism by which TLR9 may promote B cell tolerance to anti-DNA responses (32). They found that when B cells are stimulated with a complex of anti-BCR antibody and TLR9 ligand (to mimic DNA containing antigen), the B cells die in a TLR9-dependent manner, following an initial proliferative response. Interestingly, survival signals via BAFF-R or CD40 can prevent this TLR9-dependent cell death. Dependence of compromised Ars/A1 B cells on CD40 signals observed here may reflect that similar mechanisms are at play in vivo for DNA-reactive B cells. Thus, (bystander) T cell help mediated by CD40L may enable TLR9 promotion of the autoantibody response.

The relationship between anergic B cells, CD40 stimulation and T cells has been studied in several experimental systems. In the MD4 x ML5 system, alloreactive bm12 T cells can promote B cell responses and differentiation into antibody-forming cells of non-anergic MD4 B cells but not anergic MD4 x ML5 B cells (15). In vitro studies demonstrated that MD4xML5 B cells are receptive to CD40 signals but antigen receptor signaling needs to be intact with the associated upregulation of the costimulatory molecule CD86 in order for these B cells to productively respond to T cell help in vivo (15, 16, 33, 34). Instead, CD40 stimulation in the absence of BCR stimulation results in CD95 upregulation and deletion of B cells (35, 36). Our findings support the notion that autoreactive B cells are only functionally receptive to T cell help if anergy is lost and BCR signaling is restored. In line with this, we previously reported that upon SHIP-1 deletion, Ars/A1 B cells upregulate MHC class II and CD86 in response to self-antigen, suggesting their solicitation of T cell help (19). There may be situations in which provision of T cell help is sufficient to promote a response by anergic B cells. In the DNA-reactive VH3H9/VL2 model, B cells exist in an anergic state, with minimal responsiveness to BCR or anti-CD40/IL4 stimulation in vitro (37). If these B cells express influenza hemagglutinin (HA) under control of the MHC class II promoter, resulting in MHC class II presentation of HA peptide, HA-specific CD4 T cells were sufficient to drive an extrafollicular antibody response (38). Of note, in this experimental system the T cells that provide cognate help are not autoreactive, while the endogenous T cells that provide cognate help in our experimental system likely are autoreactive, possibly requiring more costimulatory signals to become activated in order to provide help. Scenarios where provision of bystander antigen (e.g. virus-derived) enables autoreactive B cells to receive help from non-autoreactive T cells have been reported to support autoreactive B cell responses (39).

Appropriate regulation of PI3K signaling is important for both central and peripheral B cell tolerance (13, 40). Suppression of PI3K pathway signaling is particularly important in peripheral tolerance mediated by anergy. Anergic B cells have elevated inositol phosphatase activity that is crucial for their unresponsive phenotype (18, 20, 41). Inducible overexpression of an inositol phosphatase induces an anergic-like phenotype in B cells (42), while (inducible) deletion of inositol phosphatases or expression of a constitutively active PI3K results in a loss of B cell anergy (19).De-repression of PI3K signaling enables BCR signaling in autoreactive B cells, providing signal 1, but, as discussed below, may have additional consequences.

The exact mechanism by which altered PI3K signaling in B cells makes them receptive to bystander T cell help is unclear. During cognate B-T cell interactions, activated antigen-specific CD4 T cells have upregulated CD40L expression, providing essential signals to antigen-specific B cells through CD40. Work from Cyster lab showed that naïve CD4 T cells express low levels of CD40L and that these expression levels remain low, partly due to continuous interactions with B cells (43). They provided evidence that these CD40-CD40L interactions can have functional consequences for B cells and showed that they provide survival signals to anergic MD4 x ML5 B cells.

CD40 ligation activates multiple signaling pathways in B cells, including the PI3K pathway. The exact pathway by which PI3K is activated downstream of CD40 in B cells remains undefined, as are the functional consequences. We hypothesize that during bystander T cell – Ars/A1 B cell interactions, Ars/A1 B cells receive CD40L/CD40 signals, including PI3K-dependent signals, that are enhanced when PI3K signaling is dysregulated, raising them above a threshold required to support an autoantibody response. At this point, we cannot exclude that other mechanisms are responsible instead. Enhanced BCR signaling has been proposed to make B cells less dependent on CD40-derived survival signals (44), possibly compensating for suboptimal CD40 signals received during bystander B-T cell interactions. Other T cell-derived signals, such as cytokines, may also play a role. Cytokine receptors also activate PI3K signaling, which may be affected by changes in inositol phosphatase activity. However, given the strict CD40L-dependence of the response of compromised Ars/A1 B cells, we suspect that the contribution of these factors will be minor.

Taken together, the work presented here suggests that dysregulation of PI3K signaling may promote loss of B cell tolerance and associated autoimmunity in several ways. Enablement of productive antigen receptor signaling is a key first step towards a loss of B cell tolerance. But increasing receptiveness to CD40L-derived signals from bystander T cells may allow activated autoreactive B cells to circumvent tolerance mechanisms imposed by T cell tolerance, raising the likelihood of a productive autoantibody response. This may have clinical relevance. In patients with autoimmune diseases, inositol phosphatase expression/activity is decreased in autoreactive B cells, which correlates with disease activity (45–47). Similarly, gain of function mutations in the PI3K subunit P110δ found in humans disrupt B cell tolerance, including B cell clones that are normally silenced by B cell anergy, leading to autoantibody production (48–51). In addition, increased expression of CD40L on lymphocytes in autoimmune patients (52) (53) and soluble CD40L in their serum (54, 55) may further promote such bystander interactions.

## Material and Methods

### Mice

Six- to sixteen-week-old mice were used in all experiments. All experiments were age-matched and sex-matched. Both genders were used in our experiments with similar results. As described previously (19), hCD20-TamCre animals (56) were intercrossed with mice carrying the rosa26-flox-STOP-YFP allele (57), generating mice in which YFP is expressed in B cells upon Cre activation. These mice were crossed with Ars/A1 B cell antigen receptor transgenic mice (21) to generate hCD20-TamCre x rosa26-flox-STOP-YFP x Ars/A1 mice. B cells from these mice will be referred to as WT Ars/A1 and are used as anergic B cell controls in adoptive transfer experiments. These mice were crossed with SHIP-1^flox/flox^ mice (donated by Dr. J. Ravetch, Rockefeller University, New York, NY) (58), PTEN^flox/flox^ mice (donated by Dr. R. Rickert, Sanford Burnham Prebys Medical Discovery Institute, La Jolla, CA) (59) or SHP-1^flox/flox^ mice (The Jackson Laboratory B6.129P2-ptpn6^tm1Rsky^/J, Strain 008336) (60) to generate mice in which SHIP-1, PTEN or SHP-1 deletion can be induced in anergic B cells respectively. In some experiments, hCD20-TamCre x SHP-1^flox/flox^ x Ars/A1 mice crossed with CD45.1 congenic C57BL/6 (The Jackson Laboratory, B6.SJL-*Ptprc^a^ Pepc^b^*/BoyJ, Strain# 002014) were used as well.

For adoptive transfer experiments, C57BL/6 mice (The Jackson Laboratory C57BL/6J, Strain # 000664), TCRα KO mice (The Jackson Laboratory B6.129S2-TCRα^tm1Mom^/J, Strain: 002116), OTII mice (The Jackson Laboratory B6.Cg-Tg(TCRαTcrb)425Cbn/J, Strain # 004194) crossed onto the TCRα KO line (OTII x TCRαKO mice) and CD8 KO mice (The Jackson Laboratory FVB.129S2(B6)-Cd8a^tm1Mak^/LcsnJ, Strain: 032563) were used as recipients.

Mice were housed and bred at the University of Colorado Denver (UCD) Anschutz Medical Campus Vivarium (Aurora, CO), with the exception of C57BL/6 mice, which were purchased from Jackson Laboratories. All experiments with mice were performed in accordance with the regulations and with approval of the University of Colorado SOM Institutional Animal Care and Use Committee.

### Adoptive transfers and tamoxifen induction

Four hours prior to adoptive transfer, recipient mice were irradiated with 200 rads to provide space for transferred cells as described previously (19). B cells from donor mice were enriched by depletion of CD43^+^ cells with anti-CD43-conjugated magnetic beads (CD43 (Ly-48), mouse; Miltenyi Biotec). In some experiments, anti-CD4-conjugated magnetic beads (CD4 (L3T4) MicroBeads, mouse; Miltenyi Biotec) were added to ensure full depletion of CD4 T cells. Resultant populations were >97% B cells based on B220 staining and FACS analysis. Donor B cells were labeled with CellTrace Violet Proliferation Kit, *for flow cytometry* (Invitrogen by Thermo Fisher) at 5μM for 3 min at RT prior to transfer in complete media (IMDM supplemented with 10% fetal calf serum (FCS), 1mM Sodium Pyruvate, 2 mM L-Glutamine, 100 U/ml Pen/Strep, 50 mg/mL gentamicin and 0.1 mM 2-Me). The labeling was stopped by adding an excess of FCS, followed by 3 washes with PBS. 0.5-2.5×10^6^ B cells in 200 μl PBS were adoptively transferred by i.v. injection. Twenty-four hours after transfer, Cre activity was induced by tamoxifen treatment. Tamoxifen (Sigma, T-5648) was dissolved in 10% ethanol (Decon Laboratories) and 90% corn oil (Sigma) at 20mg/ml. 100 μl (2mg) was injected i.p. on two consecutive days.

### In vivo antibody treatments

Blocking InVivoMab anti-mouse CD154(CD40L) antibody (Clone: MR-1; Bio X Cell) or depleting InVivoMab anti-mouse TCRbeta antibody (Clone: H57-597 (HB218); Bio X Cell) was injected i,v., or i.p. (200 μl at 1 mg/ml in sterile PBS) as indicated in experimental schema or figure legend. InVivoMab Polyclonal Armenian Hamster IgG was used for isotype control. In VivoMab anti-mouse CD40, (Clone:FGK4.5/FGK45; Bio X Cell) was injected i.v. or i.p. (200 μl at 0.25mg/ml in sterile PBS) for stimulation.

### Phenotypic analysis by staining and FACS

Single cell suspensions of splenic cells were prepared in complete media (IMDM supplemented with 10% FCS, 1mM Sodium Pyruvate, 2 mM L-Glutamine, 100 U/mL Pen/Strep, 50 mg/mL gentamicin and 0.1 mM 2-Me.), and red blood cells were lysed using Ammonium Chloride Potassium (ACK) Lysis buffer (15mM NH_4_Cl, 10 mM KHCO_3_, 0.1mM Na_2_EDTA, pH 7.4). Splenocytes were fixed and permeabilized with BD Cytofix/Cytoperm™ and stained with Dylight650-E4 anti-Ars/A1 idiotype (E4 was produced and conjugated to DyLight™ 650 NHS Ester (Fisher Scientific) in our laboratory), Dylight488 anti-GFP (anti-GFP (Goat) Antibody 600-101-215 (Rockland) conjugated to DyLight™ 488 NHS Ester (Fisher Scientific) in our laboratory), and CD138-PE-Cy7 (Clone: 281-2; Biolegend). When CD45.1 expressing Ars/A1 B cells were used in adoptive transfers, we replaced the Dylight488 anti-GFP with anti-CD45.1-FITC (Clone A20, BD Biosciences).

Events were collected on a CyAn ADP (Dakocytomation) or BD LSRFortessa™ X-20 Cell Analyzer (BD Biosciences) and analyzed using FlowJo software (Becton Dickinson & Company (BD)). In all experiments, the same gating strategy was used. Following doublet exclusion, we gated on the lymphocyte population based on FSC/SSC properties but including larger cells to capture plasmablasts, followed by gating on YFP+ E4+ events. When CD45.1 expressing Ars/A1 B cells were used as donor cells, we gated on CD45.1+ E4+ events.

### ELISPOT

For detection of IgMa Ars antibody secreting cells, microtiter plates were coated with 10 μg/ml Ars-BSA_16_ or 10 *μ*g/ml Ars-OVA_8_ in PBS. Before use, the plates were washed twice with PBS-0.05% Tween-20 and blocked with complete medium. Two-fold serial dilutions were made of splenic single cell suspensions starting at 1/50th of a spleen in the first well. The plates were incubated at 37°C at 7% CO_2_ overnight. Ars/A1-derived antibody secreting cells were detected with Biotin Mouse Anti-Mouse IgM[a] (Clone:DS-1; BD Biosciences), followed by Streptavidin-AP (SouthernBiotech). Between each step, the plates were washed 4 times with PBS-0.05% Tween-20. The plates were developed by incubating with ELISPOT development buffer (25μM 5-bromo-chloro-3-indolyl phosphate p-toluidine, 100 mM NaCl, 100 mM Tris, 10 mM MgCl2 [pH 9.5]) for 1 h. The reaction was stopped by washing the plate once with double-distilled H_2_O. The number of spots at a cell dilution in the linear range was determined, and the number of antibody-secreting cells per spleen was calculated.

### Statistics

Statistical analyses were performed using the unpaired Student’s t test. P-values <0.05 (*) were considered statistically significant. P-values <0.01 are represented by (**) and P-values <0.001 are represented by (***).

## Acknowledgements

The authors wish to thank Dr. John Cambier for his support during the early stages of this project and for valuable discussion and editing, Drs. Ross Kedl and Jared Klarquist for helpful discussion and CD8 KO mice, and Dr. Tinalyn Kupfer and the CU | AMC ImmunoMicro Flow Cytometry Shared Resource, RRID:SCR_021321, for technical assistance.

